# Applications of adeno-associated virus for 3D single-cell morphometric analysis in iPSC-derived midbrain organoids

**DOI:** 10.64898/2026.05.14.725219

**Authors:** M.B. Baeza Trallero, E. Villeneuve, P. Lépine, A.I. Krahn, C.X.Q. Chen, W. Reintsch, M.J. Castellanos-Montiel, T.M. Durcan, M.H. Berryer

**Author notes:** Corresponding authors. Correspondence to: Martin H. Berryer, Full address: McGill University, Montreal Neurological Institute, Room NWB116, 3801, University St., Montreal, QC, H3A 2B4, CANADA, Thomas M. Durcan, Full address: McGill University, Montreal Neurological Institute, Room NWB134, 3801, University St., Montreal, QC, H3A 2B4, CANADA.

## Abstract

Human midbrain organoids (hMBOs) are emerging *in vitro* models to mirror the cellular diversity and the structural complexity of the developing human brain. However, the dense neural network, limits the investigation of individual cells’ morphology or cell-cell connectivity, which is mostly restricted to fixed organoids following extensive optical clearing techniques. To better resolve individual cells within a brain organoid and for longitudinal tracking of its growth and development, we turned to adeno-associated virus (AAVs) for targeted gene delivery. In particular, we applied AAVs for expressing specific markers that provide the foundation to image individual cells within 3D hMBOs. Thus, we developed a phenotypic platform to specifically inspect the neuronal and astrocytic cytoarchitecture and to examine their connectivity in living hMBOs derived from two genetically unrelated control iPSC lines. We demonstrate that through AAV transduction, we could capture and reconstruct the 3D architecture of both neurons and astrocytes within the hMBO as a whole. Transduced cells exhibited an intrinsic heterogeneity in term of soma volume, arbor complexity and territory covered, regardless of both genetic background, age, and cell-type. Yet, these cellular morphometrics remained equivalent between the two cell lines, indicative of homogeneity in hMBO cellular development. We were able to establish longitudinal profiling of transduced cells, demonstrating how neurons and astrocytes could expand their network over time. Lastly, we describe time-lapse studies to track cellular motility and morphology fluctuations in neurons and astrocytes over time, highlighting the dynamic nature of these cells within the ramified architecture of the neural network in the developing hMBOs. Overall, our platform underscores the versatility of AAVs in studying single cell-morphometrics and cellular connectivity for longitudinal monitoring of cellular dynamics in live 3D hMBOs instead of a static snapshot.

## Introduction

Human brain organoids (hBO) are 3D multicellular self-assembling tissue derived from embryonic or induced pluripotent stem cells modelling the human brain early development (Agboola *et al*., 2021, Maisumu *et al*., 2025), recapitulating some aspects of its cellular composition (Jang *et al*., 2022; Kelley *et al*., 2022). The improvements in the organoid differentiation methodologies allow precise control of key developmental time points during the culture process to induce the formation of specific brain regions such as the cerebellum, hindbrain, forebrain, or midbrain. Timed addition of small molecules patterns the developing organoids to closely resemble the target tissue in term of cell-types, structure, and properties (Lancaster *et al*., 2013; Sloan *et al*., 2018). In addition to their potential to mirror the target tissue, patterned BOs model the complex interactions between the diverse generated neural cell-types (Quadrato *et al*., 2017; Trujillo *et al*., 2019; Velasco *et al*., 2019). Indeed, the cells produced by the developing tissue mature through close interactions with each other and their microenvironment, adopting complex cytoarchitectures and refining highly intricate cellular processes. For example, astrocytes progressively extend multiple primary processes from their soma, which further ramify in complex branching structures (Kendrick *et al*., 2017; Wang *et al*., 2025), thereby increasing individual cell volume within the developing BO. Likewise, the neurons gradually form a dense network, prolonging their axon and elaborating their dendrites into highly complex arborizations that span hundreds of micrometers within a human BO.

The development of human induced pluripotent stem cell (iPSC)–derived MBOs represent a major advance in the study of neurological disorders. Their human origin overcomes interspecies differences inherent to animal models, while preserving a 3D context that better reflects *in vivo* organization. As such, hMBOs constitute a valuable alternative model to conventional 2D cultures and animal models. Importantly, organoids generated from patient-derived iPSCs enable the development of personalized models of neurological disease. Capturing changes in cellular and structural complexity is essential to exploit hMBOs to their full potential as predictive platforms for drug discovery and therapeutic screening. Indeed, resolving cell morphology and connectivity at a single-cell level enables identifying the subtle phenotypic alterations, thereby improving the detection of disease-relevant signatures and the evaluation of candidate compounds in neurological disorders.

Investigating the elevated cellular complexity created by this experimental model requires a high level of precision to explore the full spatial structure and to extract single-cell morphology measurements. In this regard, classical histology techniques provide a solid foundation to examine tissue architecture, and spatial organization, cell type identification and quantitative cellular morphometrics; however, they require fixation and slicing of the 3D tissue, followed by immunocytochemistry with cell-specific antibodies. Inevitably, sectioning a cell spanning across hundreds of micrometers in multiple thin planes disrupts its spatial continuity, introduces distortions from tissue shrinkage and leads to fragmented views of the specific cell types of interest, making the full 3D architecture reconstruction of a neural network highly challenging. Alternatively, optical clearing approaches such as iDISCO, CUBIC, and BABB (Renier *et al*., 2014; Matsumoto *et al*., 2019; Foster *et al*., 2019) enable volumetric analysis without physical sectioning, therefore preserving the integrity of the BO. Although the whole-mount clearing methods are suitable for organoid screening (Silva Santisteban *et al*., 2017; Grenier *et al*., 2020; Rybin *et al*., 2021), these clearing techniques necessitate a chemical fixation step providing a static snapshot of the biological events and ultimately preventing real-time and longitudinal observations. Of note, the commonly used paraformaldehyde fixation method reduces antigen accessibility (Idziak *et al*., 2023; Konno *et al*., 2023) and fluorescent probes typically target specific proteins rather than uniformly labeling the entire cell at high fidelity, restricting quantitative analyses of cell shape, branching, and hindering an investigation into spatial partnership. Moreover, current clearing protocols often rely on harsh chemical treatments (methanol, tetrahydrofuran, sodium dodecyl-sulfate) reported to compromise antibody binding (Ertürk *et al*., 2012; Richardson *et al*., 2021; Tainaka *et al*., 2016).

An alternative methodology for live imaging lies in the generation of reporter iPSC lines. However, these approaches typically require extensive genome engineering procedures, making them time-consuming and technically demanding, with potential off-target effects or transcriptional silencing (Vojnits *et al*., 2022; Haideri *et al*., 2024; Puspita *et al*., 2024). Reporter expression cassettes are typically driven either by constitutive promotors (*e.g.* CAG), resulting in broad and ubiquitous labeling, or by more specific promotors that restrict expression to defined cell-types. Although some commercially available cell lines incorporate multiple cassettes/reporters, their multiplexing capacity and flexibility remain limited.

As an alternative strategy, adeno-associated virus (AAV) are potent and versatile viral vectors used for both fundamental research and preclinical studies in central nervous system gene delivery (Depla *et al*., 2020; Drouyer *et al*., 2024). AAV cell transductions are highly efficient and its long-term transgene expression often outperforms other methods for gene delivery in brain organoid (Cho *et al*., 2022). Its transduction induces low-to-no pathogenicity, in contrast to other methods, such as electroporation associated with a loss of cell viability, or lentivirus transfection related to mutagenesis risks. Moreover, the tropism of the AAVs enables precise mapping of cellular architecture and neural networks in the 3D tissue by selectively transducing the cells of interest while minimizing off-target effects (Depla *et al*., 2020; Hammond *et al*., 2017; Han *et al*., 2023). AAVs encoding fluorescent reporters (Cerulean, eGFP, tdTomato, mCherry) permit the high-resolution imaging of individual cells within the intact organoids for morphological tracing. Furthermore, the mosaic expression of the fluorescent reporter within a cell population achieved through AAV transductions in organoids facilitates segmentation and analysis at a single-cell resolution. Thus, circumventing optical clearing imaging restrictions, AAV transduction enables longitudinal and time-lapse studies for single-cell resolution tracking (Cho *et al,* 2022; Han *et al*., 2023). As a result, AAVs enable the development of an imaging protocol to assess the dynamics and spatial morphology of individual cells within a live 3D BO.

With this in mind, we developed a scalable, robust protocol to investigate cellular cytoarchitecture in live, intact hMBOs via AAVs transduction. We acquired z-stack images of the developing tissue and quantified 3D single-cell morphometrics. This workflow further highlights the relevance of AAVs in identifying, reconstructing and characterizing hSYN-tdTomato-positive neurons and gfaABC1D-eGFP-positive astrocytes. Finally, our study demonstrated the potential of AAVs in the longitudinal and dynamic investigation of individual neurons and astrocytes within these 3D BOs structures.

## Results

We developed the outlined workflow (Figure 1A), integrating AAV-mediated labeling of individual cells, combined with high content confocal imaging and 3D reconstruction to enable robust visualization and quantitative 3D cellular analysis of hMBOs. We generated hMBOs using a previously published protocol implemented in our laboratory (Mohamed *et al*., 2021). To maximize scale-up, and to minimize batch to batch variability while maintaining viability of cells in the production of the hMBOs, we used a microfabricated disk technology combined with the incubation of selective small molecules to pattern iPSC suspension towards midbrain fate 3D cellular aggregates, before their final transfer to bioreactors for maturation and long-term maintenance (Mohamed *et al*., 2022). On the day of the experiment, the hMBOs were transduced with AAVs encoding reporters under the control of the GFAP and synapsin promoters to respectively visualize astrocytes and neurons (Figure 1B). Live confocal imaging of hMBOs demonstrated that they were efficiently transduced as indicated by the expression of fluorescent reporters (Figure 1B). Acquisition of high-resolution images revealed an intricate and complex cellular network composed of multiple arborized tdTomato-positive neurons (orange cells; Figure 1B), intermingled with numerous highly branched eGFP-positive astrocytes (green cells; Figure 1B). Animations of the transduced hMBOs highlight the highly interconnected web of cell projections, as well as the uniform spatial distribution of the two cell types inside the hMBOs (Movies 1-3), demonstrating the potency of AAV vector delivery. Importantly, brightfield images confirmed that the transduction with one or a combination of AAVs did not elicit toxicity such as cell shedding or cell debris accumulation (Supplementary Figure 1A). No sign of external morphological change assessed by the area of transduced and untransduced control hMBOs was apparent over time, underscoring the preservation of the overall hMBO morphology following AAV transduction (Supplementary Figure 1B).

**Figure 1.**
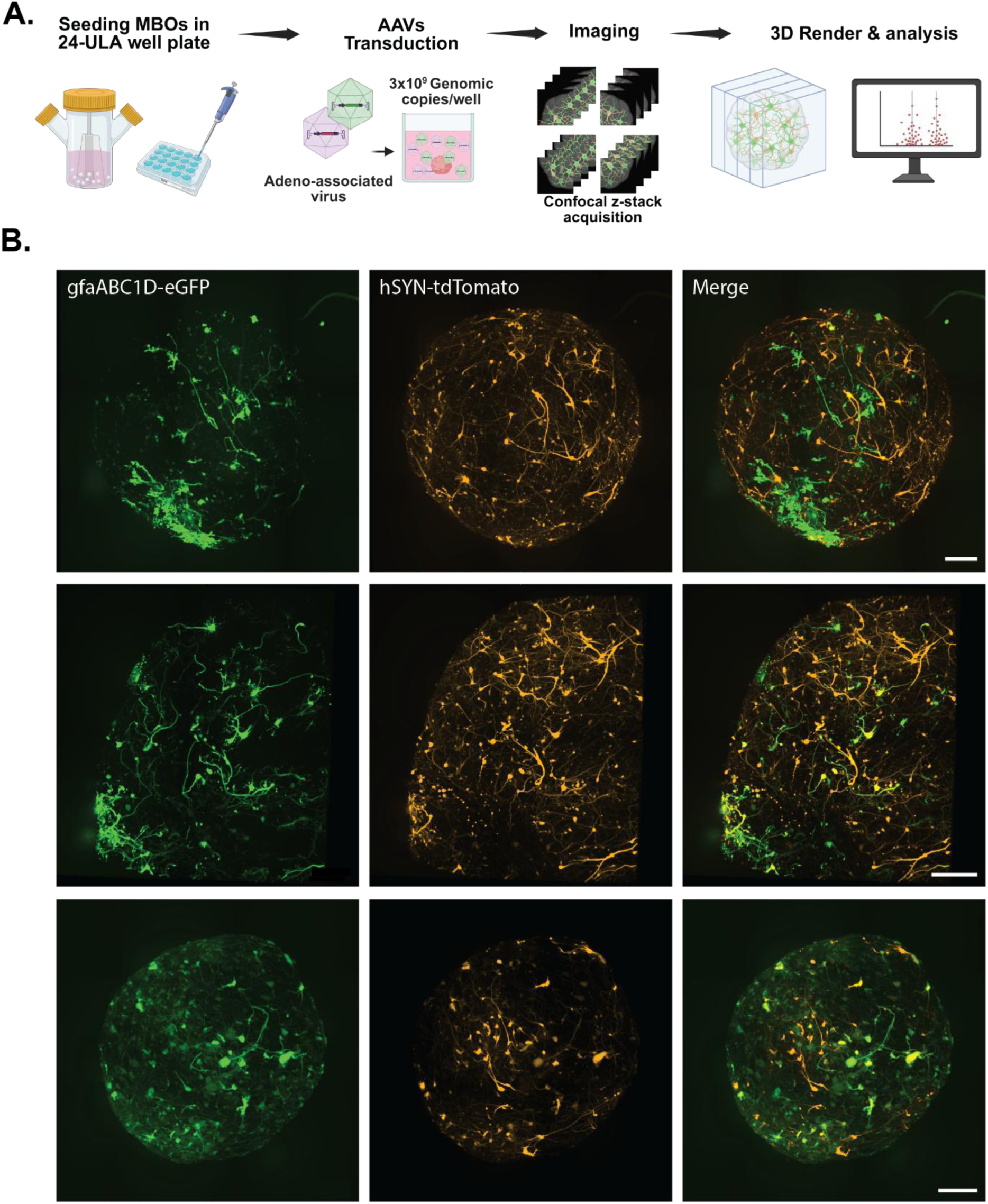
Overview of the experimental workflow for AAV-mediated labeling, confocal imaging, and 3D analysis of hMBOs. (**A**) hMBOs are seeded individually in 24-well ultra-low attachment (ULA) plates and transduced with adeno-associated viruses (AAVs) at a final concentration of 3 × 10^9^ genomic copies per well. Following incubation to allow reporter gene expression, confocal z-stack imaging is performed. Raw image stacks are subsequently converted, stitched, reconstructed, and analysed in three dimensions using IMARIS Core (version 10.2.0; RRID:SCR_007370). Image created in https://BioRender.com (RRID:SCR_018361). (**B**) Representative maximum-intensity projections of confocal stacks from three hMBOs from QPN929 (top and bottom row) and AIW002-02 (middle row) cell lines. hMBOs were transduced with AAV2/7m8-gfaABC1D-eGFP (green) and AAV2/7m8-hSyn-tdTomato (orange) and imaged 10 days after. Scale bars: 200 µm.

We noted that the eGFP expression driven by the gfaABC1D promotor was detectable as early as day 2 post-transduction, with a clear signal observed by day 4. In contrast hSYN driven tdTomato expression exhibited slower onset, becoming clearly visible around day 7 (data not shown). While reporter expression time may vary between hMBOs, both eGFP and tdTomato signals were consistently detected by day 7 to 10 post-transduction across conditions. Thus, day 10 post-transduction was chosen as our time point for image acquisition.

To validate the robustness of our method, a control line was used to generate hMBOs that were matured for 3, 6, 9, and 12 months before being incubated with the same combination of AAVs. Regardless of the maturation age, we noted consistent expression of eGFP and tdTomato exogenous proteins ten days after transduction of the hMBOs (Figure 2A), confirming the applicability of the viral approach to label both neurons (tdTomato) and astrocytes (eGFP) in our maturing hMBOs. Comparative transduction of hMBOs with a number of different capsid serotypes carrying identical reporter gene cassettes showed that all tested capsid variants successfully labelled numerous target cells (Figure 2B), demonstrating the reproducibility of this approach. Together, these findings support the broader application of AAV-based strategies to label cells of interest in hMBOs and other brain organoid subtypes.

**Figure 2.**
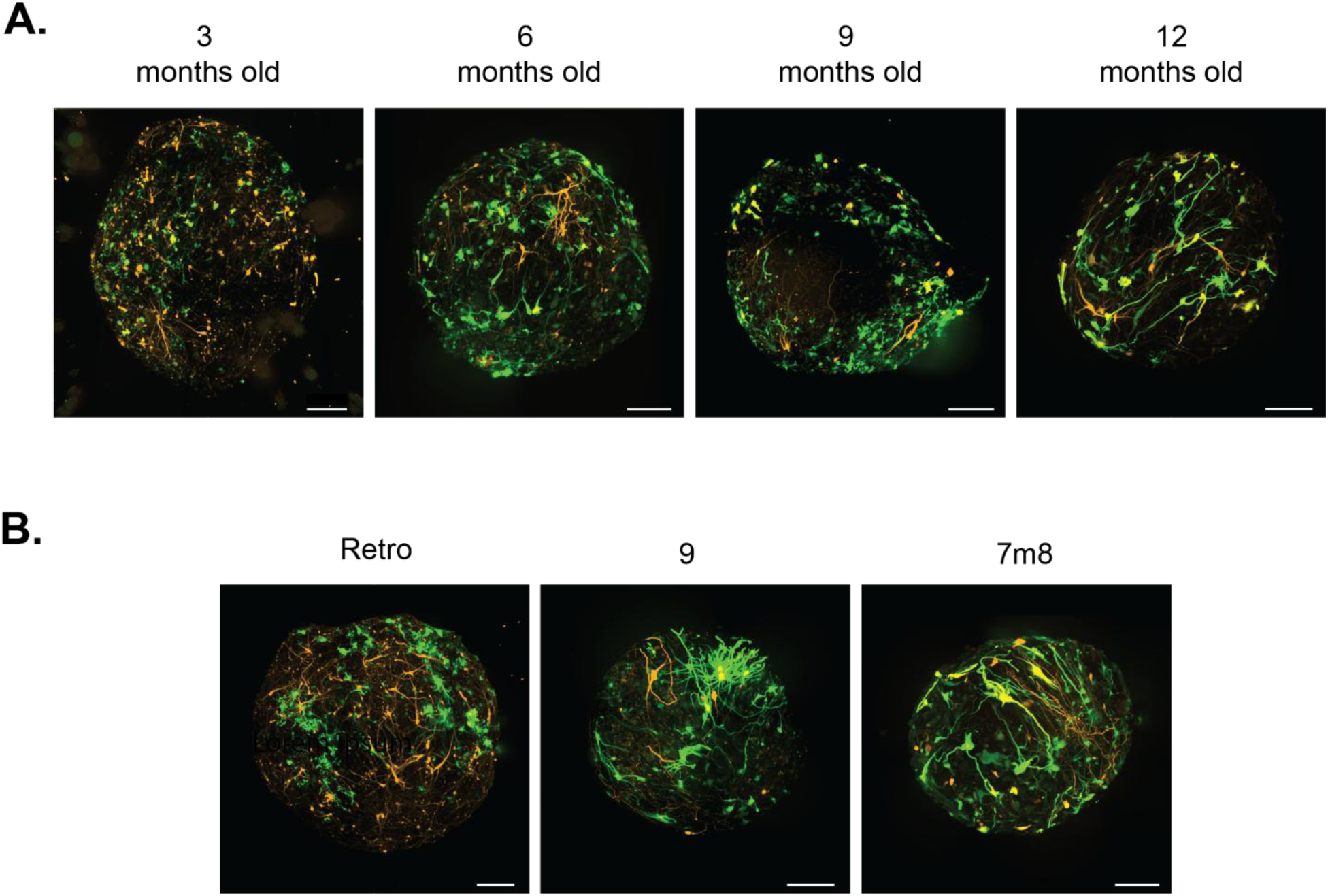
Comparison of AAV transductions. (**A**) hMBOs from QPN929 cell line transduced at 3, 6, 9, or 12 months of differentiation with AAV2/7m8-hSyn-tdTomato and AAV2/7m8-GfaBC1D-eGFP. (**B**) hMBOs from QPN929 cell line transduced with different AAV serotypes carrying hSyn-tdTomato (orange) and gfaBC1D-eGFP (green) cassettes. Serotypes (left to right): AAV2/Retro, AAV2/9, and AAV2/7m8. Images were acquired 10 days after transduction. Scale bars, 200 µm.

For the analysis of cellular morphology, confocal z-stacks images of transduced hMBOs were reconstructed. To analyse single-cell morphology within a hMBO, neuronal structures (hSYN-tdTomato) were reconstructed using the Filament Tracer module from the analysis software (Figure 3A), optimized for tracing highly arborized and elongated structures. Preserving fine morphological features, this approach enabled single-cell data extraction ranging from soma dimensions to spatial measurements evaluating the volume, sum of arbor length, number of branch points, segments, and terminal points, as well as the investigation of the cellular arborization complexity by Sholl analysis (Figure 3B-D). Quantitative analysis was performed on ten to fifteen randomly selected hSYN-tdTomato-positive neurons per hMBO, from three independent batches in either four- or six-month-old hMBOs. With hMBOs derived from two distinct cell lines, it led to 100+ virtual reconstructions per cell line. While intrinsic heterogeneity is observed in each hMBO from both lines, the soma volume andthe total length of the filament of the fully segmented hSYN-tdTomato-positive neurons was equivalent between the two cell lines, correlating with comparable numbers of segments, branch, and terminal points (Figure 3C). This analogous neuronal cytoarchitecture complexity between the two cell lines is validated by a similar arborization ramification depicted by the Sholl analysis (Figure 3D). Over 500 eGFP expressing astrocytes per cell line were segmented; with an example shown in Figure 4A. The eGFP-positive astrocytes displayed in average a comparable volume across the hMBO regardless of the cell line and maturation stage (Figure 4B), despite noticeable morphological variability among individual cells in the same 3D structure, as illustrated in the classification panel (Figure 4C). These single-cell architecture metrics uncovered by fluorescence deliver a snapshot of individual neurons and astrocytes as they grow, mature, and develop within hMBOs.

**Figure 3.**
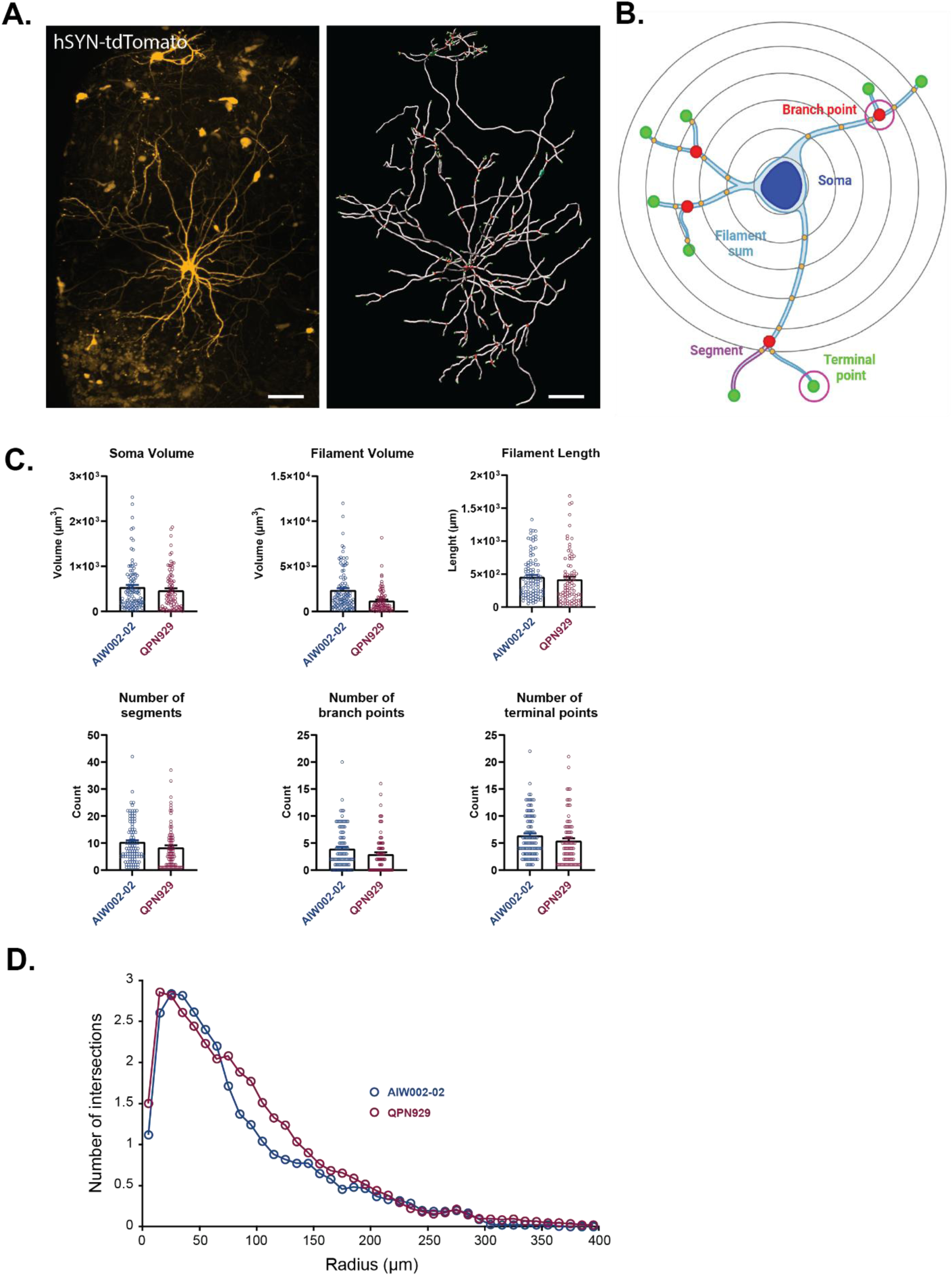
3D single-cell reconstruction and morphological analysis of tdTomato-positive neurons in hMBOs. (**A**) Left, maximum projection of transduced cells expressing tdTomato from a 10-month-old hMBO derived from QPN929 cell-line. Right, detailed reconstruction by filament tracing of labeled cells. Dendritic skeleton is shown in gray, branch points in red, and terminal points in green. Scale bars, 80 µm. (**B**) Schematic representation of neuronal morphometrics used for analysis, including soma (blue), branch points (red), terminal points (green), filaments (light blue), segments (magenta), and intersections with concentric circles (orange) for Sholl analysis. (**C**) Quantification of morphological features in tdTomato-positive neurons from 4-month-old AIW002-02 and 6-month-old QPN929. Each circle represents one reconstructed cell. Analysis was performed on 10-15 randomly selected tdTomato-positive neurons per hMBO, across three independent batches (n=3 hMBOs per batch) transduced with AAV2/7m8-hSyn-tdTomato. Data are represented as mean ± s.e.m. (**D**) Sholl analysis of tdTomato-positive cells from AIW002-02 and QPN929 hMBOs. Data corresponds to three independent batches (n = 3 hMBOs per batch, 10 neurons per hMBO). Each data point represents the average number of intersections at a given radial distance.

**Figure 4.**
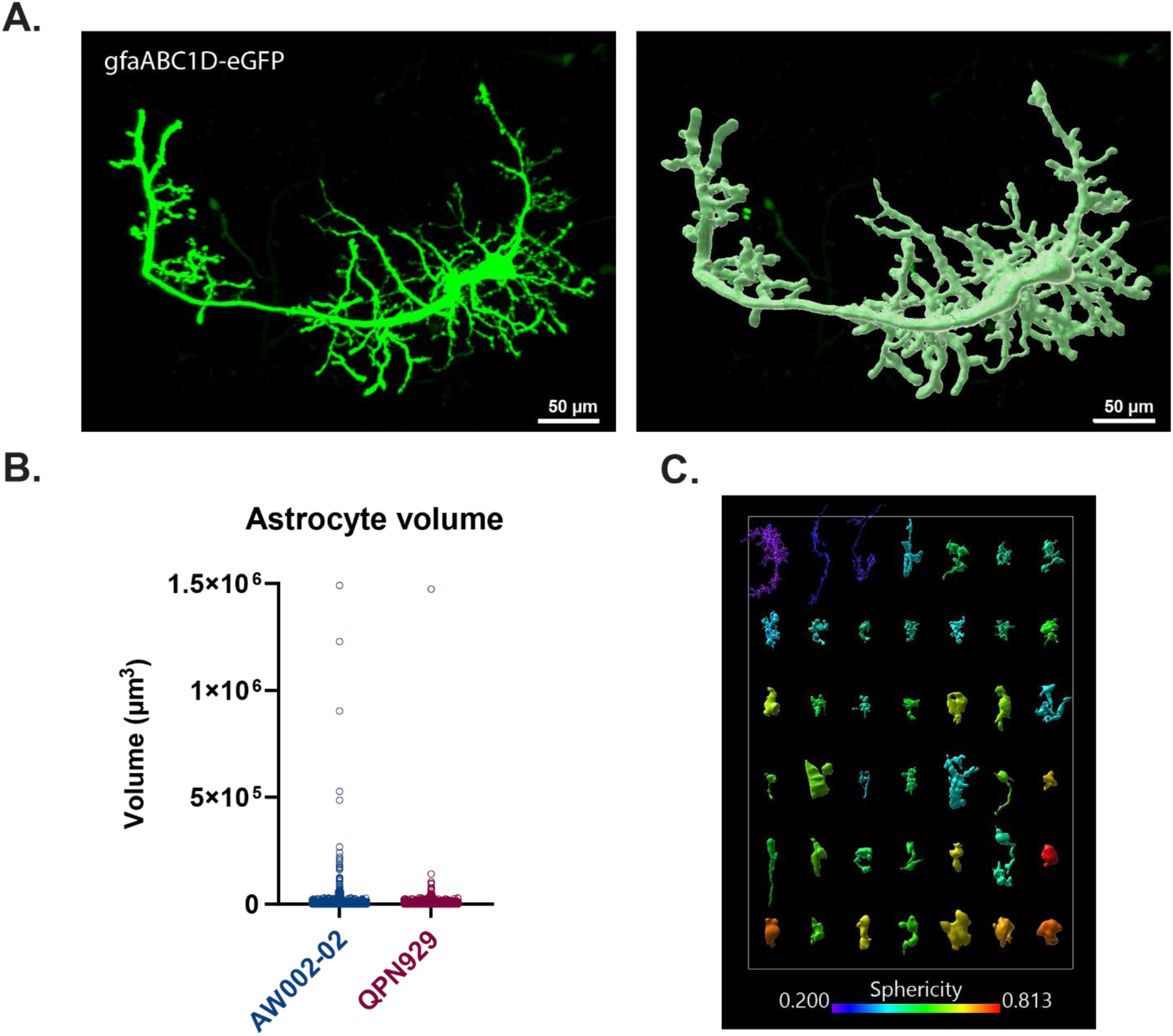
3D single-cell reconstruction and morphological analysis of gfaABC1D-eGFP-positive astrocytes in hMBOs. (**A**) Left, maximum projection of transduced astrocyte expressing eGFP from a 3-month-old hMBO derived from AIW002-02 cell-line. Right, surface-based reconstruction of the astrocyte on the left. Scale bars, 50 µm. (**B**) Quantification of volume variations across gfaABC1D-positive astrocytes from the AIW002-02 and QPN929 hMBOs. Each circle represents one reconstructed cell. Analysis includes segmented gfaABC1D-expressing cells of the hMBO, across three independent batches (n = 3 hMBOs per batch) transduced with AAV2/7m8-gfaABC1D-eGFP. Data are represented as mean ± s.e.m. (**C**) Representative panel of reconstructed individual gfaABC1D-positive astrocytes from a 3-month-old AIW002-02 hMBO. Cells sorted by volume (highest to lowest) and color-coded by sphericity. Cells are scaled to the same size for visualization purposes.

Next, we used these reconstructions to explore the cellular connectivity map in hMBOs. After hMBO transduction and segmentation of the tdTomato-positive neurons and eGFP-positive astrocytes (Figure 5A), we quantified the volume occupied by the two fluorophores. We noticed that the total volume occupied by the cells expressing the reporters represent 2 to 3% of the acquired volume of the transduced hMBOs (Figure 5B). Next, we assessed the cellular connectivity in hMBO by investigating the overlapping volume occupied by the fluorescent tags as an indicator of the cellular physical contacts. We show that approximatively 50% of the volume occupied by the fluorescent signals comes from tdTomato-positive neurons, 30% originated from the eGFP transduced astrocytes, and 20% were shared by the two fluorophores (Figure 5C).

**Figure 5.**
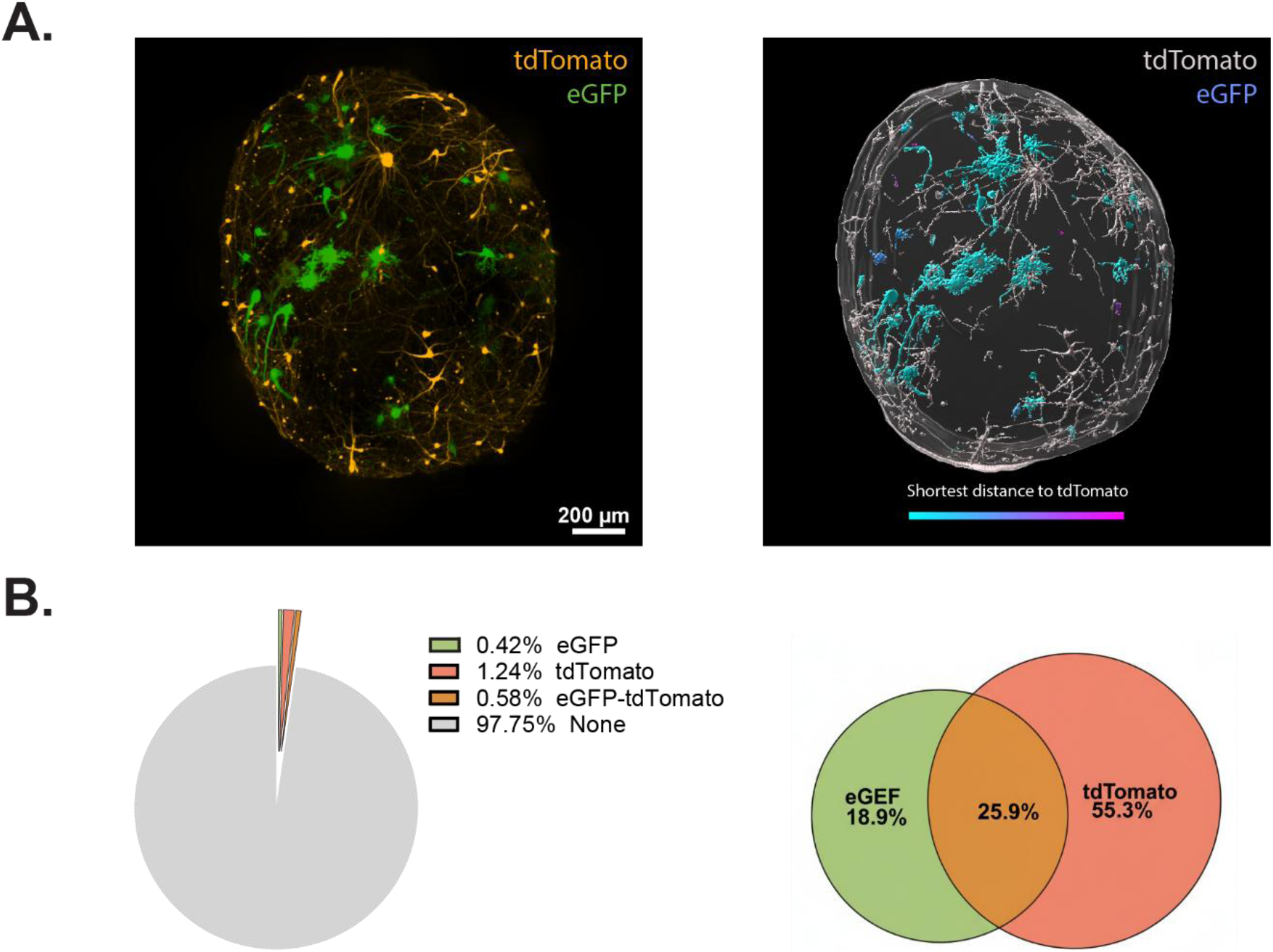
Cellular connectivity analysis in hMBO. (**A**) Left, representative maximum-intensity projections from a hMBO imaged 10 days after transduction with both AAVs hSYN-tdTomato and gfaABC1D-eGFP. Right, surface-based reconstruction of the same hMBO showing the whole surface of the hMBO, hSyn-tdTomato in gray and gfaABC1D-eGFP surface color-coded according to the shortest distance to hSyn-tdTomato surface from closest (cyan) to furthest (pink). (**B**) Left, pie chart showing the proportion of total hMBO volume occupied by eGFP, tdTomato, and their overlap. Right, Venn diagram representing the relative contribution of each fluorescent signal and their overlap within the labeled volume.

Importantly, we noticed that the expression of the marker persisted up to one-month post-transduction, with well-defined intensity signals in both channels and low-to-no detectable cytotoxicity (Figure 6A), supporting the basis for longitudinal studies. We transduced a total of twelve hMBOs in two independent batches, with either hSYN-tdTomato or gfaABC1D-eGFP, or a combination of both AAVs. Acquired with the same laser power and exposure time across the entire image acquisition procedure for comparisons across the experiment, we observed an expansion of the surface covered by neurons or astrocytes expressing their fluorescent marker over time (Figure 6A), demonstrating how, as the hMBOs mature, so does the density of the neural network. Correlating with those observations, the volume occupied by the fluorescent expressing cells in each hMBO gradually rose (Figure 6B), whereas the intensity reached saturation at later time points, making accurate quantification challenging. Through this approach, we successfully tracked and quantified the morphology fluctuations of the exact same eGFP-positive astrocytes. The resulting 3D renderings revealed the constant remodelling of astrocytic architecture and the ability of the workflow to capture changes in cell morphology over time (Supplementary Figure 2).

**Figure 6.**
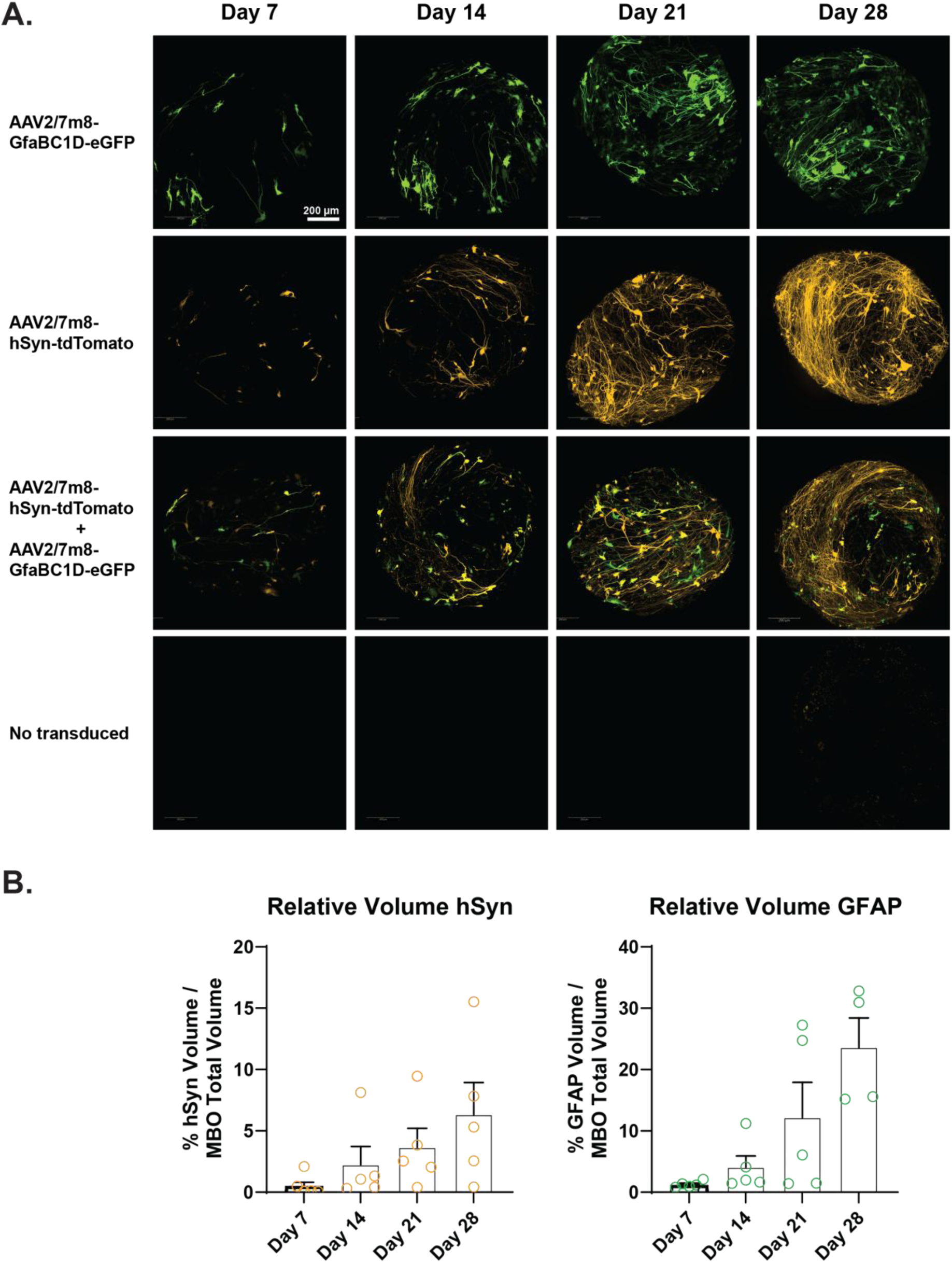
Persistent AAV-mediated fluorescent reporter expression in 13-month-old QPN929 hMBOs. (**A**) Representative maximum projections from the same four hMBOs, acquired 7, 14, 21, and 28 days post-transduction. hMBOs were transduced with either AAV2/7m8-gfaBC1D-eGFP (top), AAV2/7m8-hSyn-tdTomato (2^nd^ row), both (3^rd^ row), or untransduced (bottom). Scale bar, 200 µm. (**B**) Quantification of the volume occupied by either the tdTomato or eGFP tags divided by the acquired volume of the hMBO 7, 14, 21, and 28 days post-transduction. Data are represented as mean ± s.e.m.

Finally, we performed time-lapse imaging to assess live readouts from the eGFP expressing astrocytes in 3D hMBOs. One week after AAVs transduction, images were acquired automatically and sequentially over 72 hours, without noticeable phototoxicity (Figure 7A). Using a segmentation algorithm from the analysis software, we tracked the dynamics of the transduced cells in the living tissue (movies 4-7). Astrocytes were observed to dynamically move and change shape, with corresponding fluctuations in different features as cell volume or displacement length over time (Figure 7 and Movies 4-7). These changes were quantified at both the population and single-cell level (Figure 7B). These observations highlight the value of time-lapse imaging for capturing dynamic cellular behaviors in 3D systems. Altogether, our approach provides a robust framework for real-time and longitudinal analysis of cellular morphology and dynamic in 3D cellular structures without the need for tissue dissociation or terminal procedure.

**Figure 7.**
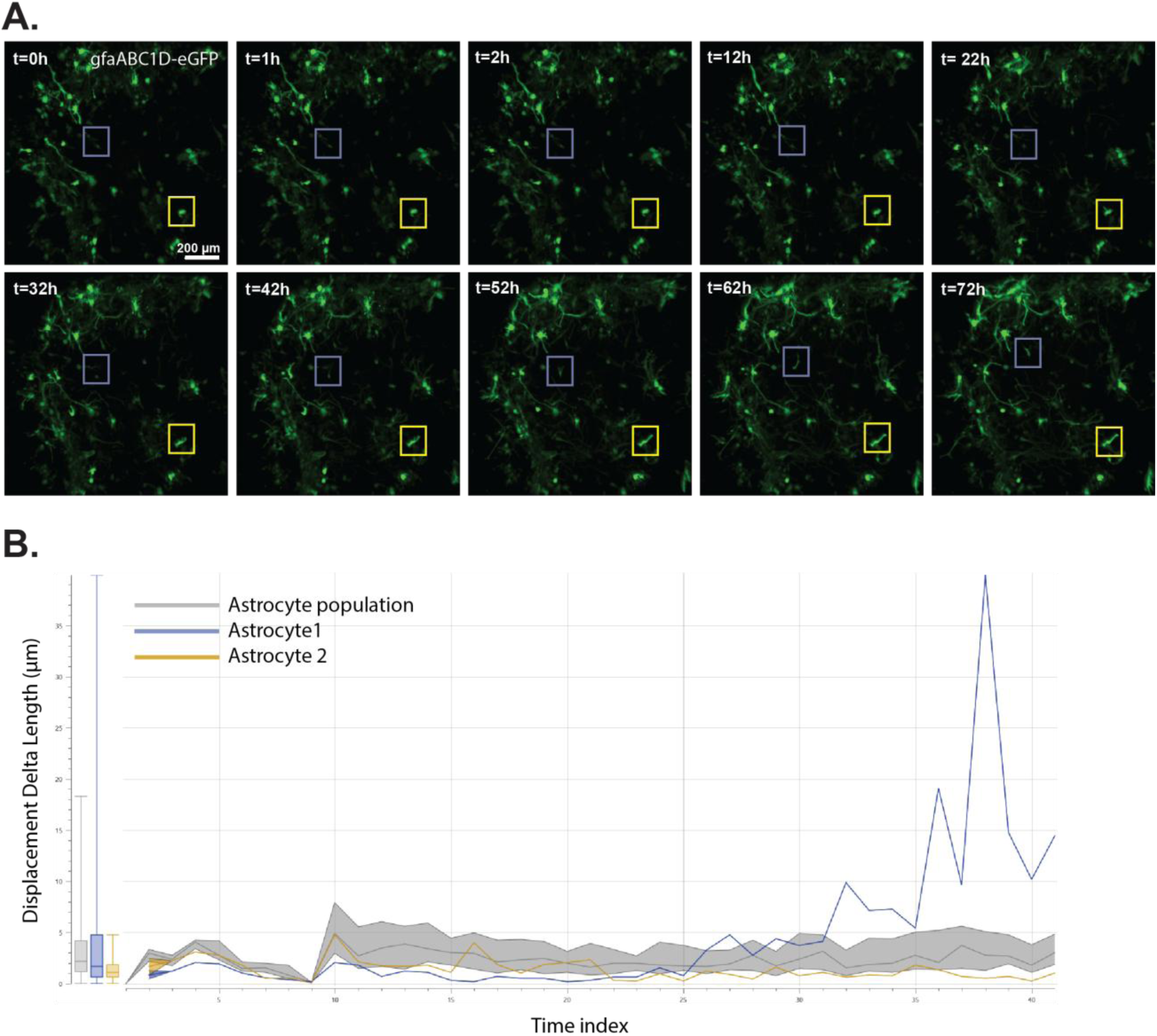
Time lapse imaging of transduced hMBOs. (**A**) Representative maximum projections from a transduced hMBO (AAV2/Retro-gfaBC1D-eGFP) acquired 7 days post transduction. The same region of the hMBO was acquired at 41 time points over 72 hours. Individual gfaABC1D-eGFP-positive cells randomly selected for analysis at single-cell level are outlined by squares (blue and yellow). Scale bars, 200 µm. (**B**) Representation of frame-to-frame displacement over time (µm) for the hMBO shown in (A). Population-level data are shown in gray (mean ± SD); trajectories of two representative single cells highlighted in (A) are shown in blue and yellow, respectively.

## Discussion

In this study, we developed a versatile, robust platform based on AAVs transduction that enables high-resolution, single-cell morphology measurements in live hMBOs. By combining cell-type specific fluorescent labelling, 3D confocal image acquisition and computational reconstruction, we provide a pipeline that allows to track cells in live tissue. Importantly, this strategy enables longitudinal studies of cellular architecture shifting the analysis from static snapshots to more dynamic studies.

A central strength of the method lies in the use of AAVs as a highly efficient and minimally to non-invasive tool to label a substantial proportion of the tissue of interest. Across multiple serotypes tested, we observed robust and reproducible transductions, consistent with the tropism and the stability of transgene expression (Hagerty *et al*., 2019; Abulimiti *et al*., 2021). In contrast to alternative delivery methods such as electroporation or lentivirus transfection, which are often associated with reduced viability and risk of genomic alterations respectively (Cho *et al*., 2022), AAVs provide stable reporter gene expression with limited toxicity, making them suitable for long term studies. In our system, sustained reporters’ expression over several weeks establishes a strong foundation for extended analysis and facilitates the tracking of cellular morphology and spatial relationships over time.

An additional strength of this workflow lies in its compatibility with time-lapse imaging, enabling repeated visualization of the same hMBO regions over time. This approach allows direct tracking of individual cells and their morphological evolution within an intact 3D environment. In contrast to endpoint-based analysis, this approach captures dynamic processes such as changes in cell volume, structural complexity, and spatial relationships as they occur. Importantly, because imaging is performed without disrupting tissue integrity, these observations remain anchored in their native spatial context, enhancing the physiological relevance of the data.

The use of cell-type specific promotors, including hSYN and gfaABC1D, driving distinct fluorescent reporters, enables simultaneous labeling of two major neural populations within the same hMBOs, neurons and astrocytes.This dual-labeling strategy provides significant advantages by allowing both the detailed characterization of individual cell morphologies and the visualization of interactions between distinct cell types. Moreover, AAV transduction in organoids enables mosaic labeling within a specific cell population, facilitating accurate single-cell segmentation. By preserving the structural integrity of the tissue and avoiding the disruptive processes, our workflow captures these interactions in a context that is closer to physiological conditions.

To extract quantitative information from these complex datasets, we implemented a reconstruction pipeline that combines filament tracing for neuronal structures, and surface-based reconstruction for astrocytes. This approach reflects the intrinsic morphological differences between these cell-types, with neurons exhibiting highly arborized processes and astrocytes occupying spatially defined territorial domains and underscores the adaptability of the software to measure different features. While these tools were selected to optimize segmentation accuracy and feature extraction of each population, both approaches remain adaptable depending on experimental needs or the desired output.

Beyond structural characterization, our workflow facilitates the quantification of cell-to-cell spatial relationships. Metrics derived from surface overlaps provide a basis for estimating the cellular interactions and exploring the spatial organization of neural networks in 3D tissues. By transforming imaging data into quantitative descriptors, this methodology makes possible comparative analysis across experimental conditions, hMBO developmental stages, cellular genetic backgrounds, or AAV serotypes.

Despite these advances, some limitations must be considered. First, imaging depth is restricted to approximatively 200µm from the hMBO surface, potentially biasing analyses to peripheral regions and overlooking deeper tissue compartments. Second, slight variations in AAV transduction efficiency can influence the subsequent identification and, reconstruction of the labeled cells. Third, while our segmentation pipeline enables detailed reconstructions, it relies on thresholding, user-defined parameters, that may introduce variability, highlighting the need for further standardization or automation. Finally, although the promotors used here provide a relatively high degree of cell specificity, they may not be entirely exclusive as occasional double-positive cells were observed.

Looking forward, this pipeline opens multiple options for further research. Its compatibility with live imaging makes it particularly attractive for studying disease phenotypes in patient derived hMBOs, as well as for assessing the impact of drugs or other perturbators in real time. While developed for hMBOs, this approach is also adaptable to other types of organoids and to conventional 2D cultures, broadening its applicability across experimental approaches. Further developments could include multiplexed AAV strategies using additional promotor-reporter combinations to label more than two cell types at once (*e.g.* oligodendrocytes, neural progenitors). This type of multiplexed methodology would allow to capture complex interactions between different cell populations and provide a more complete view on how cells are organized and behave within the living and developing tissue. In addition, the use of this workflow combined with complementary approaches such as calcium imaging, machine learning segmentation algorithms, or conditional genetic cassettes could further enhance its functional and analytical potential. Finally, improvements in imaging technologies including light-sheet microscopy in association with advanced clearing strategies may help overcome current limitations in tissue penetration and imaging resolution.

## Material and methods

### Ethical approval

All human materials used were approved by the McGill University Health Centre Ethics Board under project number 22-03-02 (IRB Study Number A03-M19-22A) for induced pluripotent stem cells. All methods were performed in accordance with the relevant guidelines and regulations.

### Human iPSC cultures

Two control cell lines generated at the Montreal Neuronal Institute were used for this study: (*i*) AIW002-02 (CBIGi001-A, also known as IPSC_0063, https://hpscreg.eu/cell-line/CBIGi001-A) reprogrammed from peripheral blood mononuclear cells from a consent informed 37 years old Caucasian male (Chen *et al*., 2021); (*ii*) QPN929 (CBIGi004-A, also known as IPSC_0034, https://hpscreg.eu/cell-line/CBIGi004-A) reprogrammed form a consent informed 50 years old Caucasian female. Briefly, iPSCs were cultured and expanded on plates coated with Matrigel® hESC-qualified Matrix in mTeSR medium. Cells were maintained at 37°C with 5% CO_2_ with daily media changes and passaged when reaching 75-85% confluency by incubation in Gentle Cell Dissociation medium for 5 minutes at 37°C to obtain a single-cell suspension before replating in mTeSR medium supplemented with the ROCK inhibitor Y27632. Mycoplasma assays performed with MycoAlert™ detection kits were run on a weekly basis. For clarity, providers and catalog numbers for the medium, small molecules, and kit are listed in Table 1.

**Table 1.**

| Item | Provider | Catalog number |
| --- | --- | --- |
| Matrigel hESC qualified matrix | Corning | 354277 |
| mTeSR medium | ThermoFisher Scientific | 05851/05852 |
| Genlte Cell Dissociation medium | StemCell Technologies | 7174 |
| Y27632 | Selleck Chemicals | S1049 |
| MycoAlert detection kit | Lonza | LT07-318 |
| Accutase | StemCell Technologies | 7920 |
| DMEM/F12 | Gibco | 10565-018 |
| GlutaMAX | Gibco | 35050-061 |
| B27 without vitamin A | Gibco | 12587-010 |
| MEM-NEAA | Gibco | 11140-035 |
| 2-mercaptoethanol | Millipore Sigma | 8057400005 |
| Heparin | Sigma | H3149-100KU |
| SB431542 | Selleck Chemicals | S1067 |
| Noggin | Preprotech | 120-10C |
| CHIR99021 | Selleck Chemicals | S2924 |
| Trypan Blue Stain | Gibco | 15250-061 |
| 948-well EB disk | eNUVIO | eN-eb948u-002 |
| FGF8 | Preprotech | 100-25 |
| SHH | Preprotech | 100-45 |
| Neurobasal medium | Life Technologies | 21103-049 |
| BDNF | Preprotech | 450-020 |
| GDNF | Preprotech | 450-010 |
| Ascorbic acid | Millipore Sigma | A5960 |
| db-cAMP | Carbosynth | ND07996 |
| Laminin | Sigma | L2020 |

### Generation of human iPSC-derived hMBOs

hMBOs were generated from the two previously described iPSC cell lines using published methodologies implemented in our laboratory (Mohamed *et al*., 2021; Mohamed *et al*., 2022). Briefly, cells from a passage number <15, were passaged at least twice after thawing; iPSC colonies were dissociated with Accutase for 5 min at 37°C, then quenched with an equal volume of DMEM/F-12+ Antibiotic-Antimycotic. Cells in suspension were pelleted for 3 min at 1200 rpm and resuspended to single-cell in neuronal induction medium (for 50 mL: 47.286 mL DMEM/F12 + Antibiotic-Antimycotic /Neurobasal (1:1), 0.5mL GlutaMAX^TM^-I (1%), 1 mL B27 without vitamin A (1:50), 0.5 mL N2 (1:100), 0.5 mL MEM-NEAA (1%), 0.175 µL 2-mercaptoethanol, 50 µL Heparin (1µg.mL-1), 50 µL SB431542 (10µM), 50 µL Noggin (200ng.mL-1), 50 µL Y27632 (10µM), 13.5 µL CHIR99021 (0.8 µM)), counted with Trypan Blue to evaluate cell viability before uniformly dispensing 10×10^6^ cells on top of a 948-well ULA-coated EB DISK (eNUVIO). Dishes were then centrifuged at 1200 rpm for 10 minutes, kept in an incubator at 37 °C, 5 % CO_2_. Two days later, neural induction medium was delicately changed to neural induction medium without ROCK inhibitor and four days after seeding the neural induction medium was swapped for midbrain patterning medium (for 50 mL: 49.925 mL of neural induction medium without ROCK inhibitor supplemented with 50 µL FGF8 (100 ng.mL-1) and 25 µL SHH (100 ng.mL-1)) for the next three days followed by undiluted Matrigel® hESC-qualified Matrix embedding and a successive incubation of 30 minutes at 37 °C, 5 % CO_2_. The midbrain patterning medium was then replaced for the tissue induction medium (for 50 mL: 47.354 mL Neurobasal medium, 0.5mL GlutaMAX^TM^ (1%), 1 mL B27 without vitamin A (1:50), 0.5 mL N2 (1:100), 0.5 mL MEM-NEAA (1%), 0.175 µL 2-mercaptoethanol, 50 µL FGF8 (100 ng.mL-1), 25 µL SHH (100 ng.mL-1), 12.5 µL insulin (2.5 µg.mL-1), 8.5 µL laminin (200 ng.mL-1), 50 µL Pen/Strep) for the next 24 hours before finally transferring the embedded 3D cellular aggregates in final differentiation medium (for 50 mL: 47.363 mL Neurobasal medium, 0.5 mL GlutaMAX^TM^ (1%), 1 mL B27 without vitamin A (1:50), 0.5 mL N2 (1:100), 0.5 mL MEM-NEAA (1%), 25 µL BDNF (10 ng.mL-1), 25 µL GDNF (10 ng.mL-1), 25 µL ascorbic acid (100 µM), 12.5 µL db-cAMP (125 µM), 50 µL Pen/Strep) into 500 mL capacity spinner flask (Corning, Cat# 4500-500) placed on top of a magnetic stirrer set with a constant agitation of 42 rpm in an incubator at 37 °C, 5 % CO_2_. Medium was changed once a week and regularly tested for mycoplasma contamination with MycoAlert^TM^ detection kits. For clarity, providers and catalog numbers for the medium, small molecules and kit are listed in Table 1.

### Adeno-associated virus transduction

AAVs were produced and kindly provided by the COVF (Canadian Optogenetics and Vectorology Foundry, Quebec, CANADA, RRID:SCR_016477). Upon reception, cryovial units of AAVs comprising several capsid serotype variants (9, Retro, 7m8) were aliquoted and stored at −80 °C. Each vector carried a single fluorescent reporter cassette driven by a cell-specific promoter, either gfaABC1D-eGFP (SKU: 1744-aav) or hSYN-tdTomato (SKU: 919-aav). Unless otherwise indicated, experiments were performed using the 7m8 serotype. Prior to transduction, hMBOs were transferred from bioreactors to 24-well ultra-low attachment plates (Costar®, 3473), with one hMBO per well. hMBOs were then transduced with AAVs diluted in culture medium to a final concentration of 3×10^9^ genomic copies per hMBO. Transduced hMBOs were kept under constant agitation on an orbital shaker (ThermoScientific Cat # 88-881-103) at 100 rpm in an incubator at 37 °C, 5 % CO_2_. Half of the medium was changed every 72 hours until the start of the experimental procedure.

### Live confocal imaging acquisition

On the day of the experiment, the living transduced hMBOs were moved to black 96-well plates (Falcon, Cat# 353219), better suited for imaging purposes. z-stacks images covering a total depth of 200µm inside the living hMBOs were acquired. Images were acquired on an OperaPhenix Plus High-Content Screening System (Revvity, ex PerkinElmer), environmental control of 37 °C and 5 % CO2, using a 20x water-immersion objective (NA=1.0). Imaging was performed across 9 to 16 fields with 10% overlap, capturing 100 z-stacks at 2µm step sizes for detailed imaging. Three channels were acquired: Brightfield, the 488-laser line (Excitation: 488 nm Emission: 515-550 nm) and the 568-laser channel (Excitation: 561 nm; Emission: 570-630 nm). For longitudinal and time-lapse recordings, acquisition parameters were set up at the start of the studies and kept unchanged all along the procedures to ensure the best comparison (generating sometimes intensity saturation at later time points). For time-lapse protocol, sequential images from the three channels were acquired every 15 minutes up to 2 hours then every 2 hours for the next 70 hours.

### 3D reconstruction anwd image analysis

Raw z-stack images from adjacent acquired fields were converted to *.ims format, stitched, and rendered by a visualization and analysis software, IMARIS 10.2.0 (Oxford Instruments) for visualization of the cellular architecture across large regions of a hMBO. 3D rendering, segmentation, and measurements were performed using the following modules: Core (3D/4D rendering and visualization), Surfaces (astrocyte segmentation and volumetric analysis), Filament Tracer (neuronal reconstruction and branching analysis), MeasurementPro (quantitative feature extraction), Vantage (data visualization and statistical comparison), and Batch (automated protocol execution). Surface and filament reconstructions were performed semi-automatically using the Batch processing module to ensure consistent parameter settings across all samples within the same experiment. Reconstructions and segmentations were subsequently inspected and manually corrected, when necessary, without blinding.

### Data collection and analysis

For the morphometric analysis, all gfaABC1D-eGFP-positive cells and 10-15 hSYN-tdTomato transduced neurons per hMBO were quantified, for a minimum of 9 hMBOs per cell line distributed in 3 independent batches of at least 3 hMBOs. Regarding the longitudinal analysis, we transduced a total of 14 QPN929 human midbrain hMBOs distributed in 2 independent batches. Images for time-lapse recording experiments were acquired from 2 hMBOs from 2 independent batches. Quantitative cellular morphometrics were extracted, imported and plotted on Prism (GraphPad, version 11.0; RRID:SCR_002798). Transduction efficiency were assessed as per 3D volume covered by fluorescence divided by total volume of the hMBO acquired. Rather than comparing the different readouts between cell lines, or between time-points for potential statistical significance, this study was intended to illustrate the range of possible readouts that could be investigated. Therefore, no statistical comparison, no test for outliers, no test for normality and no sample calculation were performed. Illustrations were realized with Adobe Illustrator (RRID:SCR_010279) and Microsoft PowerPoint (RRID:SCR_023631).

## Author contribution

CXQC generated and maintained the human iPSC lines; PL produced the hMBOs; PL and MBBT maintained the hMBOs; MBBT, MHB, and EV transduced the hMBOs; MBBT, MHB, PL, and AIK acquired the images; AIK and WER assisted for image acquisition set up; MJCM set up IMARIS software in the laboratory and helped with pipelines on this software. MBBT, EV, and MHB reconstructed the data avatars; MBBT and MHB analysed the data and made the figures; MHB, MBBT, and TMD developed the workplan and experimental studies for the manuscript. MHB, MBBT, and TMD wrote the manuscript and edited it. All listed authors contributed with their edits to the manuscript.

## Abbreviations & Nomenclature

3D: three-dimensional
AAV: adeno-associated virus
BABB: benzyl alcohol and benzyl benzoate
BDNF: brain-derived neurotrophic factor
BO: brain organoid
CUBIC: Clear, Unobstructed Brain/Body Imaging Cocktails and Computational analysis
eGFP: enhanced green fluorescent protein
GDNF: glial-derived neurotrophic factor
GFAP: glial fibrillary acidic protein
hMBO: human midbrain organoid
hSYN: human synapsin
iDISCO: immunolabeling-enabled three-dimensional imaging of solvent-cleared organs
iPSC: induced pluripotent stem cells
Pen/Strep: Penicillin-Streptomycin
RRID: Research Resource Identifier (see scicrunch.org)
ULA: ultra-low attachment

## Acknowledgements

TMD received funding for this project from the Translational Initiative in De-Risking Neurotherapeutics (TRIDENT), and the New Frontiers in Research Fund (NFRF). Additional funding was provided to TMD through a Platform support grant from Brain Canada for the Canadian Optogenetics and Vectorology Foundry (RRID: SCR_016477). We acknowledge the “Canadian Foundation for Infrastructure (CFI) led John R. Evans Leaders Fund (JELF)” for supporting purchase of Opera Phenix. Thanks to Marie-Eve Paquet and the COVF team for kindly providing AAVs and related technical assistance. Thanks to María José Castellanos-Montiel for her valuable training and guidance on the use IMARIS software and 3D reconstruction workflows.

## Data Availability

The data that support the findings of this study are available from the corresponding author upon reasonable request.

## Figure Captions

**Supplementary Figure 1.** Brightfield images of hMBOs at different time points after transduction. (**A**) Representative brightfield images of individual 13-month-old QPN929 hMBOs transduced with either AAV2/7m8-hSyn-tdTomato (top), AAV2/7m8-gfaABC1D-eGFP (2^nd^ row), both (3^rd^ row), or untransduced (bottom) at days 7, 14, 21, and 28 in culture after transduction. Scale bar, 500µm. (**B**) Quantification of the surface of transduced hMBOs on day 7, 14, 21, and 28 post-transductions. Data are represented as mean ± s.e.m.

**Supplementary Figure 2.** Single-cell longitudinal volumetric analysis in hMBO. (**A**) Representative maximum intensity projections and surface-based reconstructions of gfaABC1D-eGFP-positive cells (fluorescence in green; reconstructed cells colored by object ID) tracked in a 13-month-old QPN929 hMBO, acquired at days 7, 11, 14, and 28 (from top to bottom) post-transduction with AAV2/7m8-gfaABC1D-eGFP. Scale bar, 100 µm and 50 µm (right column). (**B**) Quantification of the volume fluctuations across time from the reconstructed cells in (A).

**Supplementary movies 1-3** Representative 3D reconstruction of the 3 hMBOs in Figure 1. hMBOs were transduced with AAV2/7m8-gfaABC1D-eGFP (green) and AAV2/7m8-hSyn-tdTomato (orange) and imaged 10 days after.

Movie 1. 3D Reconstruction Hmbo1.mp4

Movie 2. 3D Reconstruction Hmbo2.mp4

Movie 3. 3D Reconstruction Hmbo3.mp4

**Supplementary movie 4:** Representative time lapse movie of a transduced hMBO and tracking of a group of cells within it. The hMBO was transduced with AAV2/Retro-GfaBC1D-eGFP (green) then acquired 7 days post transduction. The same region of the hMBO was acquired at 41 time points over 72 hours. <u>Movie 4. Time lapse on hMBO.mp4</u>

**Supplementary movie 5-7**: Over time analysis at single-cell level of 2 individual gfaABC1D-eGFP-positive cells randomly selected (blue and yellow) within the organoid in Supplementary movie 4. Movies 5 and 6 illustrate morphological and position changes over time within the organoid. Surface-based reconstruction of the cells is shown in blue and yellow. Scale bar, 20 µm. Movie 7 plots comparative changes in morphology and displacement (speed, µm/s) of the two cells previously mentioned over time. Reconstructed cells are color coded by sphericity at every time point. Scale bar, 100 µm

Movie 5. Astrocyte 1 (blue).mp4

Movie 6. Astrocyte 1 (yellow).mp4

Movie 7. Speed-sphericity tracks for astrocytes 1 and 2.mp4

