## Supplementary figures and images for "Applications of adeno-associated virus for 3D single-cell morphometric analysis in iPSC-derived midbrain organoids"

### Supplementary Figures

#### Supplementary Figure 1

A.

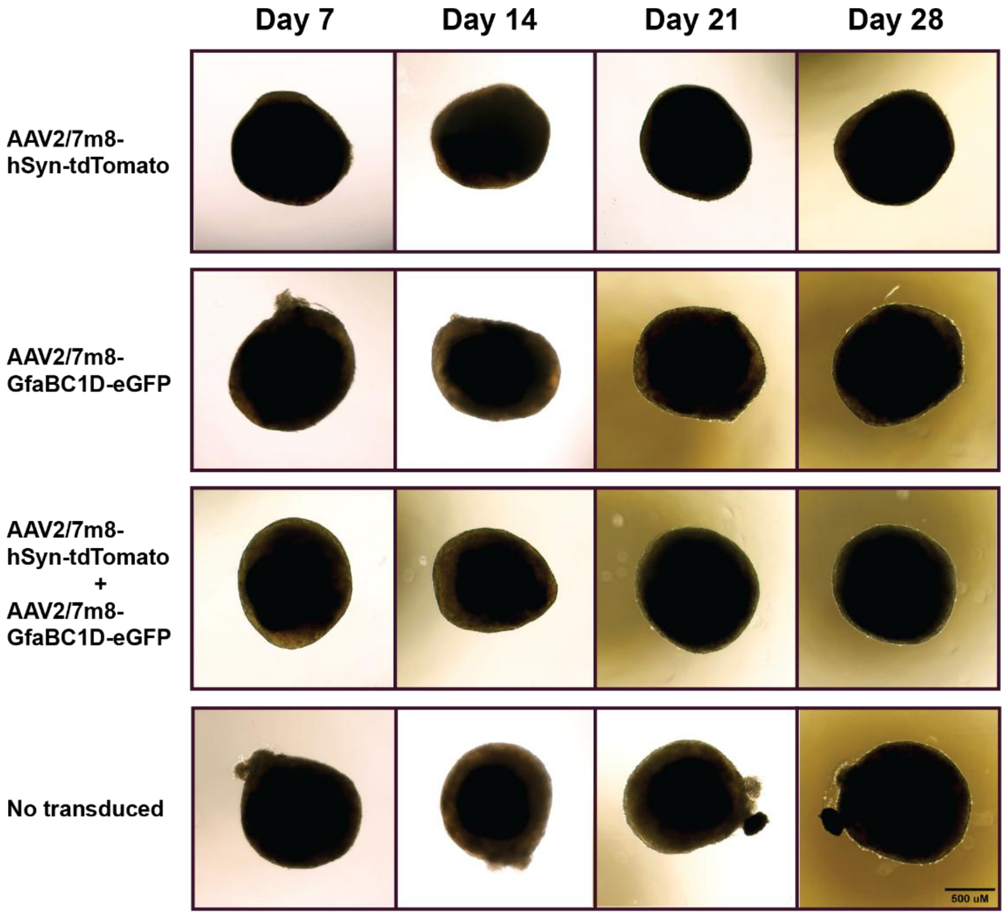


B.

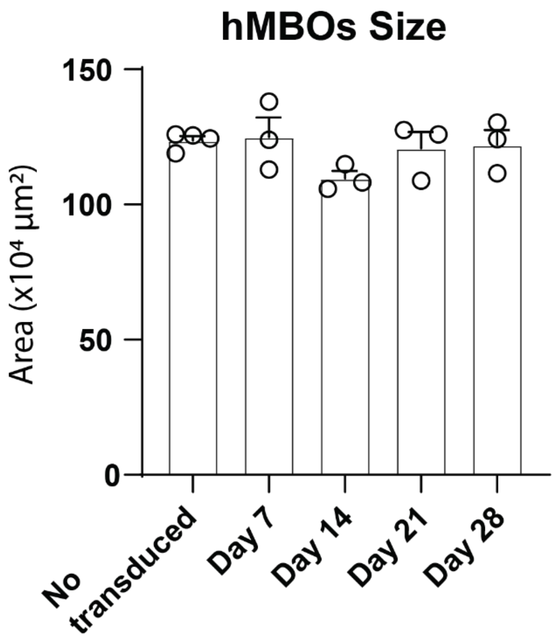

##### Supplementary Figure 2

A.

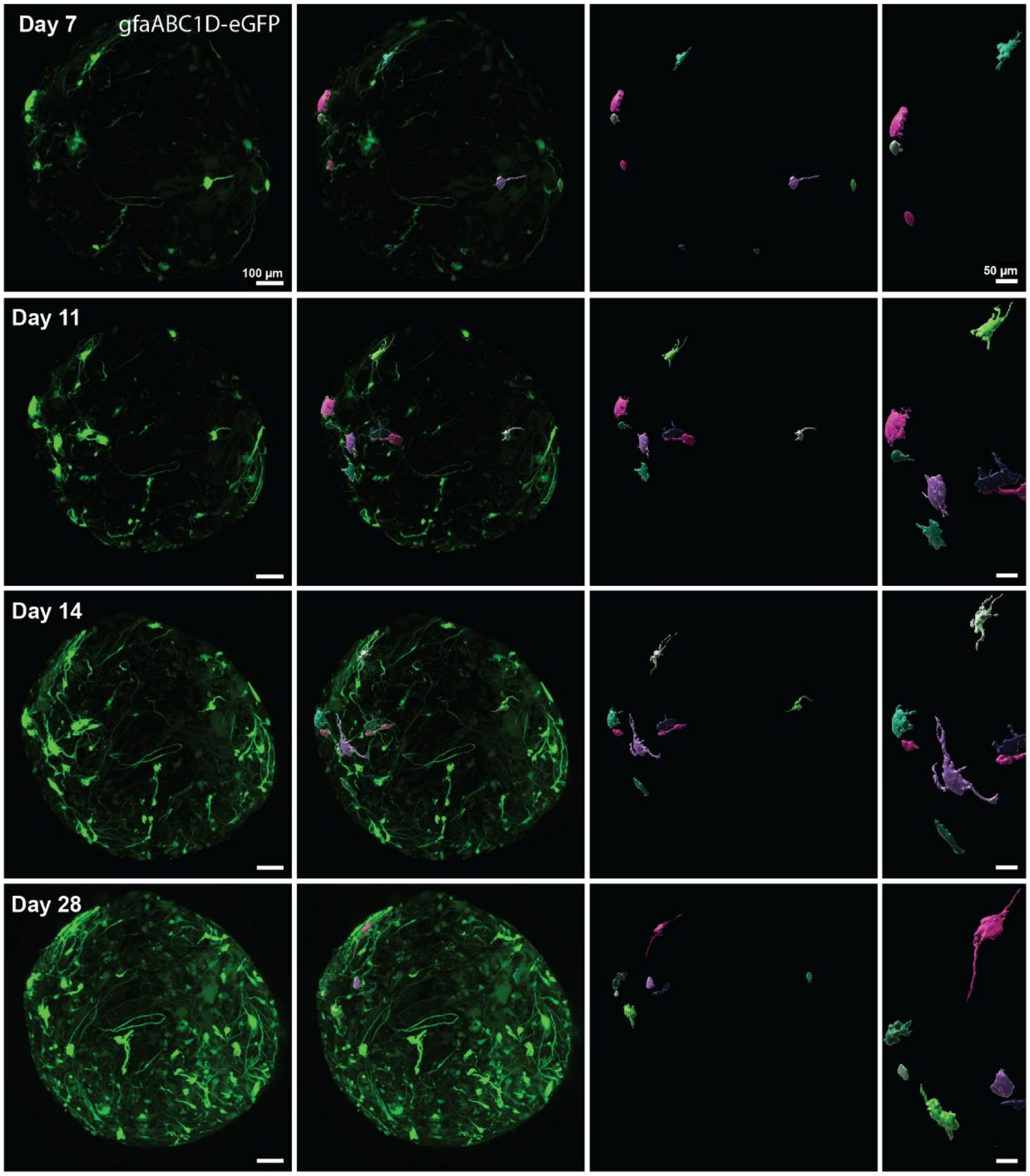


B.

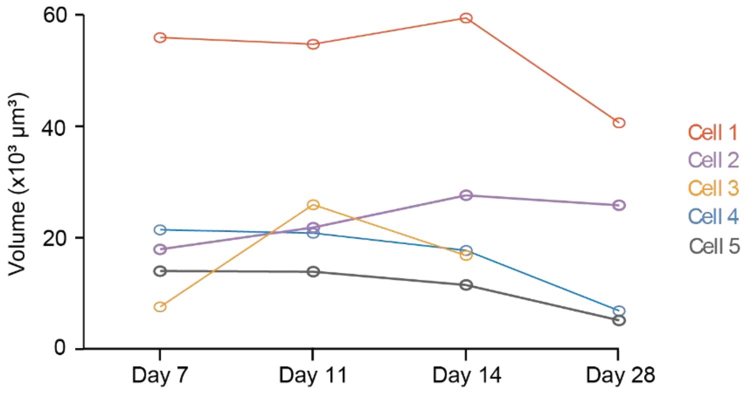
